# Investigation of the Antimicrobial Activity of Soy Peptides by Developing a High Throughput Drug Screening Assay

**DOI:** 10.1101/045294

**Authors:** Rekha Dhayakaran, Suresh Neethirajan, Xuan Weng

## Abstract

**Background:** Antimicrobial resistance is a great concern in the medical community, as well as food industry. Soy peptides were tested against bacterial biofilms for their antimicrobial activity. A high throughput drug screening assay was developed using microfluidic technology, RAMAN spectroscopy, and optical microscopy for rapid screening of antimicrobials and rapid identification of pathogens.

**Methods:** Synthesized PGTAVFK and IKAFKEATKVDKVVVLWTA soy peptides were tested against *Pseudomonas aeruginosa* and *Listeria monocytogenes* using a microdilution assay. Microfluidic technology in combination with Surface Enhanced RAMAN Spectroscopy (SERS) and optical microscopy was used for rapid screening of soy peptides, pathogen identification, and to visualize the impact of selected peptides.

**Results:** The PGTAVFK peptide did not significantly affect *P. aeruginosa*, although it had an inhibitory effect on *L. monocytogenes* above a concentration of 625 µM. IKAFKEATKVDKVVVLWTA was effective against both *P. aeruginosa* and *L. monocytogenes* above a concentration of 37.2 µM. High throughput drug screening assays were able to reduce the screening and bacterial detection time to 4 h. SERS spectra was used to distinguish the two bacterial species.

**Conclusions:** PGTAVFK and IKAFKEATKVDKVVVLWTA soy peptides showed antimicrobial activity against *P. aeruginosa* and *L. monocytogenes*. Development of high throughput assays could streamline the drug screening and bacterial detection process. *General significance:* The results of this study show that the antimicrobial properties, biocompatibility, and biodegradability of soy peptides could possibly make them an alternative to the ineffective antimicrobials and antibiotics currently used in the food and medical fields. High throughput drug screening assays could help hasten pre-clinical trials in the medical field.

## 1. Introduction

Antibiotic resistance is a significant challenge throughout the world today [43]. Extensive or ineffective use of antibiotics and antimicrobials in both the medical and food industry has led to increases in antibiotic resistance [43]. In nature, microorganisms do not always live in their planktonic forms, rather, they are found in dynamic complex structures called biofilms [18, 46]. The adaptability of pathogenic bacterial biofilms to different physiological conditions and treatments enables them to emerge as resistant strains [13]. Therefore, there is a need to develop new antimicrobial agents and antibiotics that could effectively destroy pathogenic biofilms. Bioactive antimicrobial peptides (AMPs) are a new class of bio-pharmaceuticals released from proteins that exhibit unique antimicrobial, antioxidant, and antihypertensive properties [10, 19, 29, 36]. These are specific protein fragments encrypted in the amino acid sequences, which have a positive impact on health and bodily functions [19, 20, 25]. By binding to specific receptors on target cells, they can regulate functional properties. For example, *lactoferricin* possesses antiviral, antimicrobial, antifungal, anti-inflammatory, and immuno-regulatory properties, while c*aseniate* possesses antioxidant, antihypertensive, and antimicrobial properties [4, 11, 40]. AMPs have some common features, including short chain sequences (generally 12-50 amino acid sequences), strong cationic characters (net charge ranging from +2 to +9), heat stability, and amphipathic natures, while they can be derived from plants, animals, insects, and engineered microorganisms, or could be synthesized [24, 35]. Although peptides are diverse in the amino acid sequences that make up their structure, they can be classified into four main categories: (a) α-helices, (b) β-sheets, (c) extended helices, and (d) loop-forming structures [35]. Currently, numerous efforts are focussed towards understanding the antimicrobial efficacy and mode of action of AMPs on bacterial cells.

AMPs interact with bacterial cell membranes and penetrate cells changing the pH gradient, membrane potential, and osmotic regulation, thereby affecting respiration [24]. Influencing intra-cellular mechanisms, such as immuno-modulatory relationships, disrupting the cell membrane and forming pores are other possible modes of action of AMPs [27]. Pore formation is an important mechanism by which peptides cause cell death. Four models have been used to explain this phenomenon: *(i) the toroidal model* where the peptides that attach to cell membranes aggregate, and the lipid monolayers bend continuously acquiring positive curvature. Through this pore, the peptides enter and destroy the cell’s contents. In *(ii) the carpet model*, AMPs cover the surface of the cell like a carpet until a threshold is reached to form membrane patches in which the lipids aggregate. This weakens the bilayer and forms pores. Therefore, membrane disruption occurs leading to lysis. In *(iii) the barrel-stave model*, bundles of peptides oligomerize and form membrane pores on the microbe, thus enabling them to interact with the hydrophobic core. Therefore microbes die either by membrane disruption, cell leakage, or loss of polarization. In the final *(iv) aggregate channel model*, peptides penetrate bacterial cells and form clusters and aggregates. The water molecules then facilitate the leakage of ions from the cell [24]. AMPs are also implicated in the inhibition of DNA, protein, and RNA synthesis, interference with cytokinesis, alteration of cytoplasmic membrane formation, and the inhibition of enzymatic activities [24].

Peptides usually occur as α-helix or β-sheets or as a combination of both. There are two common characteristics shared by most AMPs, a net positive charge and amphipathic structure. These two factors help peptides bind the cell membrane, increasing permeability [26]. It is generally accepted that long chain peptides insert themselves into bacterial cells forming pores that lead to cell death. When the net positive charge of a peptide is high, it has the ability to interact effectively with the negatively charged bacterial outer membrane [24]. The length and composition of amino acid sequences affect the formation of aggregates and dissociation during the interaction [26]. Therefore, peptides may hold promise as antimicrobial agents, although screening methodologies for testing new antimicrobials are lacking.

Conventional assays are widely used in testing and screening the effects of antimicrobial agents and antibiotics. These protocols are of use in the fields of food safety, pharmacology, environmental, and water quality monitoring, although there are some disadvantages, such as time consumption, intensive labor, the requirement of large amounts of sample or reagents, and the exposure to contamination [22, 41]. In addition, conventional assays cannot be used for intermediate testing, such as examining the kinetics of microbes, exploring topography, or characterizing the biochemical spectra. The other major issue with conventional assays is that the actual environment of microbial growth cannot be mimicked in flasks, test tubes, or microtitre wells [22]. Microfluidics technology is a fast-growing field that overcomes the disadvantages of these conventional assays by being able to provide high throughput screening, real-time examination of samples, usage of very low amounts of reagents, rapid screening, parallel testing of multiple drugs, all while saving space [9, 21, 33, 42, 44, 45, 48]. High throughput screening (HTS) platforms have improved drug delivery processes and are playing a vital role in the discovery of anti-cancer drugs and the characterization of drug metabolism and cytotoxicity [1, 23, 37, 49].

In this study, the antimicrobial efficacy of soy based antimicrobial peptides (PGTAVFK and IKAFKEATKVDKVVVLWTA) against *Listeria monocytogenes* and *Pseudomonas aeruginosa* was examined initially using conventional microbiological assays. A microfluidic high throughput drug screening assay was then developed to detect and screen the effect of the antimicrobial peptide IKAFKEATKVDKVVVLWTA in conjunction with RAMAN spectroscopy and optical microscopy to identify the pathogens and view the possible mode of action on bacterial cells.

## 2. Materials and Methods

### 2.1. Bacterial Strains

*P. aeruginosa* strain PA-76, isolated from canine ear skin infections was obtained as a gift from the Ontario Veterinary College (OVC), Guelph, Canada. *L. monocytogenes* strain C379, an isolate from chicken wieners was obtained as a gift from the Canadian Research Institute for Food Safety (CRIFS), University of Guelph, Canada.

### 2.2. Culture conditions

*P. aeruginosa* and *L. monocytogenes* were obtained as frozen stocks (-80°C). From this, the bacterial strains were streaked onto 5% sheep blood agar plates (SBP) and incubated at 37°C for 24 h. For subsequent experiments, colonies from these plates were used for culturing bacteria in Mueller Hinton-II broth (MHB) for *P. aeruginosa*, and brain heart infusion broth (BHI) for *L. monocytogenes*.

### 2.3. Peptide Synthesis

Synthesized soy peptides PGTAVFK and IKAFKEATKVDKVVVLWTA were purchased from Genemed Synthesis Inc., (Texas, USA). The molecular weight of the first short chain peptide PGTAVFK was 718.4 g/mol, and the other long chain peptide IKAFKEATKVDKVVVLWTA was 2146.4 g/mol. They were synthesized by 9-fluorenylmethoxicarbonyl (fmoc) solid-phase synthesis and purified to 98.62% and 97.59%, respectively using a C-18 high performance liquid chromatography (HPLC) reverse column [14]. Upon receiving the peptides, they were stored at 4°C. A 7 mg amount of PGTAVFK peptide was dissolved in 1 ml of sterile phosphate buffered saline (PBS; pH = 7.0) and stored at −25°C until further use. A 10 mg of IKAFKEATKVDKVVVLWTA peptide was dissolved in 1 ml of sterile PBS and stored at −25°C until further use. The stock concentrations were therefore 5 mM for PGTAVFK and 4.66 mM for of IKAFKEATKVDKVVVLWTA.

### 2.4. Antimicrobial Assay

The microdilution assay was adapted from Motyl *et al* [30]. *P. aeruginosa* was grown in 6 ml cation adjusted Mueller Hinton broth (Sigma Aldrich, Canada), and *L. monocytogenes* was grown in 6 ml Brain Heart Infusion Broth (Thermo Scientific, Canada). Mueller Hinton broth is the commonly used media for performing susceptibility tests for fast growing clinically isolated bacteria. While performing preliminary experiments, it was found that *L. monocytogenes* did not grow in Mueller Hinton broth. BHI is rich in nutrients and is suitable for fastidious organisms like *L. monocytogenes*. Both bacteria were grown at 37°C in the respective broth until the cell density was above the 0.5 McFarland barium sulfate standard, while the cells are in the logarithmic growth phase. This inoculum was then diluted 1:140 times in the respective media. 100 µl of media was added to each well tested in a 96-wells microtitre plate. Then, 200 µl of 5 mM PGTAVFK and 2.3 mM IKAFKEATKVDKVVVLWTA soy peptide solutions were added to the first well (column 1), and each row was serially diluted two-fold to column 11 of the 96-well microtitre plate. Column 12 was reserved for the control and did not contain any peptide solution. Tests were separately carried out with the two peptides. Lactoferricin was used as positive control, as it is a better-characterized peptide that has been shown to have antimicrobial activity against *L. monocytogenes* and *P. aeruginosa* [32, 34]. Concentrations of lactoferricin ranging from 8 µg/ml to 0.1 mg/ml have been shown to be effective against these organisms; our results were consistent with these observations (data not shown). Using a pipette, 100 µl of the diluted inoculum was added to all wells in the plate. A final inoculum of 4-8 × 10^5^ CFU/ml was achieved for each bacterial species. The final soy peptide concentrations ranged from 2.5 mM in column 1 to 2.44 µM in column 11. Plates were then covered and incubated at 37°C. At 24 h, the plates were taken out, and the optical density was read using a Biotrak II plate reader (Amersham Biosciences, Canada). All assays were performed in triplicate.

### 2.5. Microfluidic Platform

The microfluidic device was designed using AutoCAD 2015 (Autodesk Inc, USA). The vision behind designing this device was to fabricate a 3D high throughput drug screening device consisting of a glass bottom layer, a polydimethylsiloxane (PDMS) second layer with a bacterial dispenser, loading channels, and 24 incubation chambers, and a final top PDMS layer with 4 concentration generating gradients and connective channels. The glass base was 70 mm X 70 mm; the central reservoir for dispensing bacteria was designed to have a diameter of 2.5 mm. The 24 incubating chambers were 2 mm in diameter, and the loading channels were 12.75 mm long and 60 µm wide. The 4 concentration gradient generators (width = 100 µm) consisted of 8 inlets (each was 0.75 mm in diameter) to flow different concentrations of soy peptide with PBS buffer. The gradient generators were uniformly designed to ensure that they use laminar flow and diffusive mixing to form a steady mixture of PBS and peptide solution. The inlets were only present on the top layer to allow for proper mixing and the flow of peptide solution. The connecting channels were about 10-15 mm in length, while the width was varied from 100 to 30 µm to avoid the backflow of bacteria into these channels. Once the device was designed, the mask was custom made from Fine Line Imaging (Colorado, USA). From the mask, master moulds were fabricated using photo-lithographic technique. The design of this 3D high throughput microfluidic platform is shown in Figure 1.

A total of 35 g of PDMS elastomer base was mixed with 3.5 g of curing agent (Dow Corning Sylgard 184, USA). It was then poured out on the silicon wafer individually. The wafers were then desiccated and baked at 60°C for the PDMS mould to harden. The device was then removed from the wafer, cut, and holes were punched (Harris Uni-Core, Sigma Chemical Co., USA). The inlet holes on the top layer were 0.75 mm in diameter, the bacterial dispensing chamber was 0.75 mm in diameter, and the 24 incubating chambers were 2 mm in diameter. Both the layers along with the glass base were plasma cleaned (Harrick Plasma, Ithaca, USA) to remove debris and to enhance the bonding between various layers. Then, the layers were assembled and baked at 60°C for 30 minutes to strengthen the bond.

**Figure 1.**
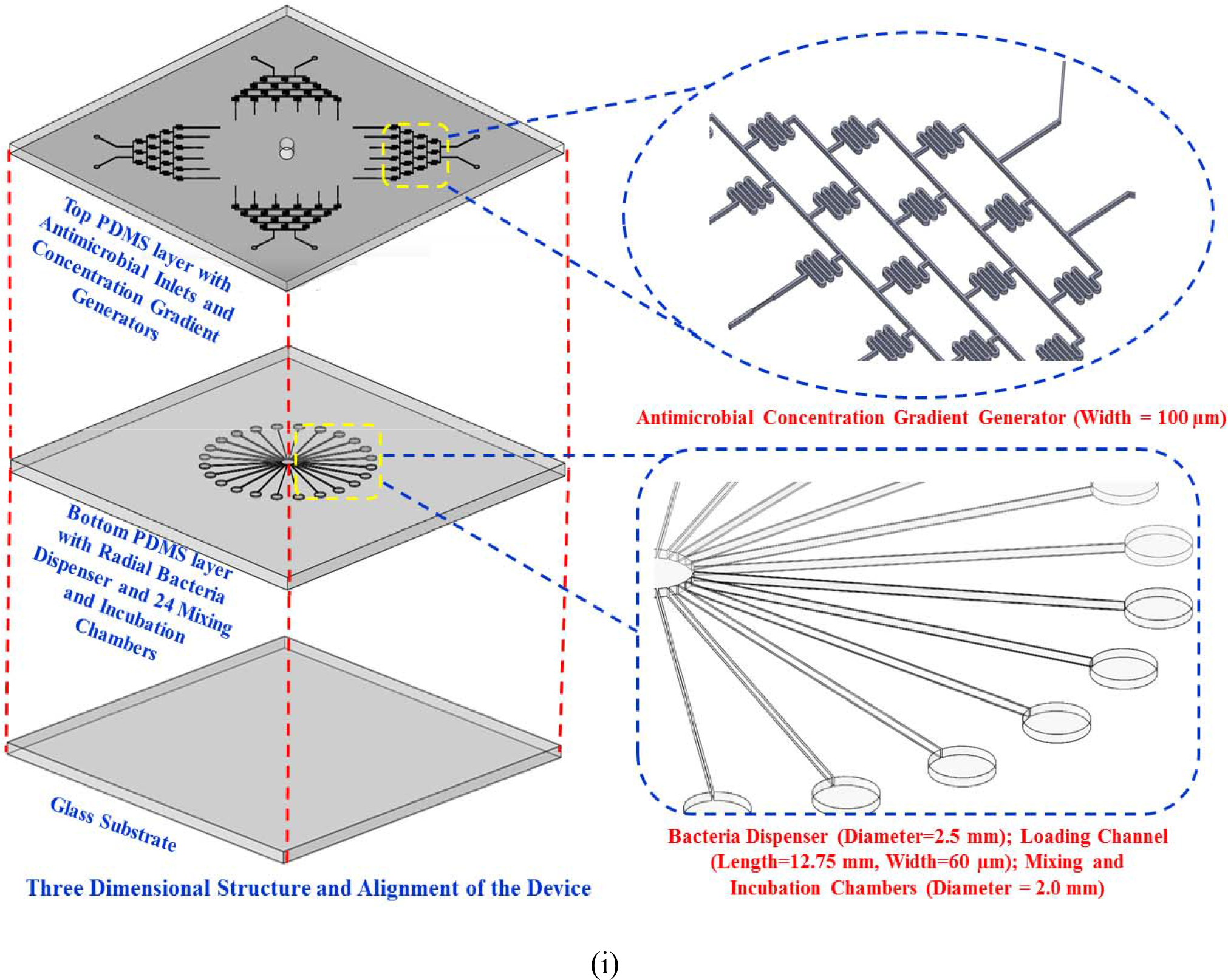
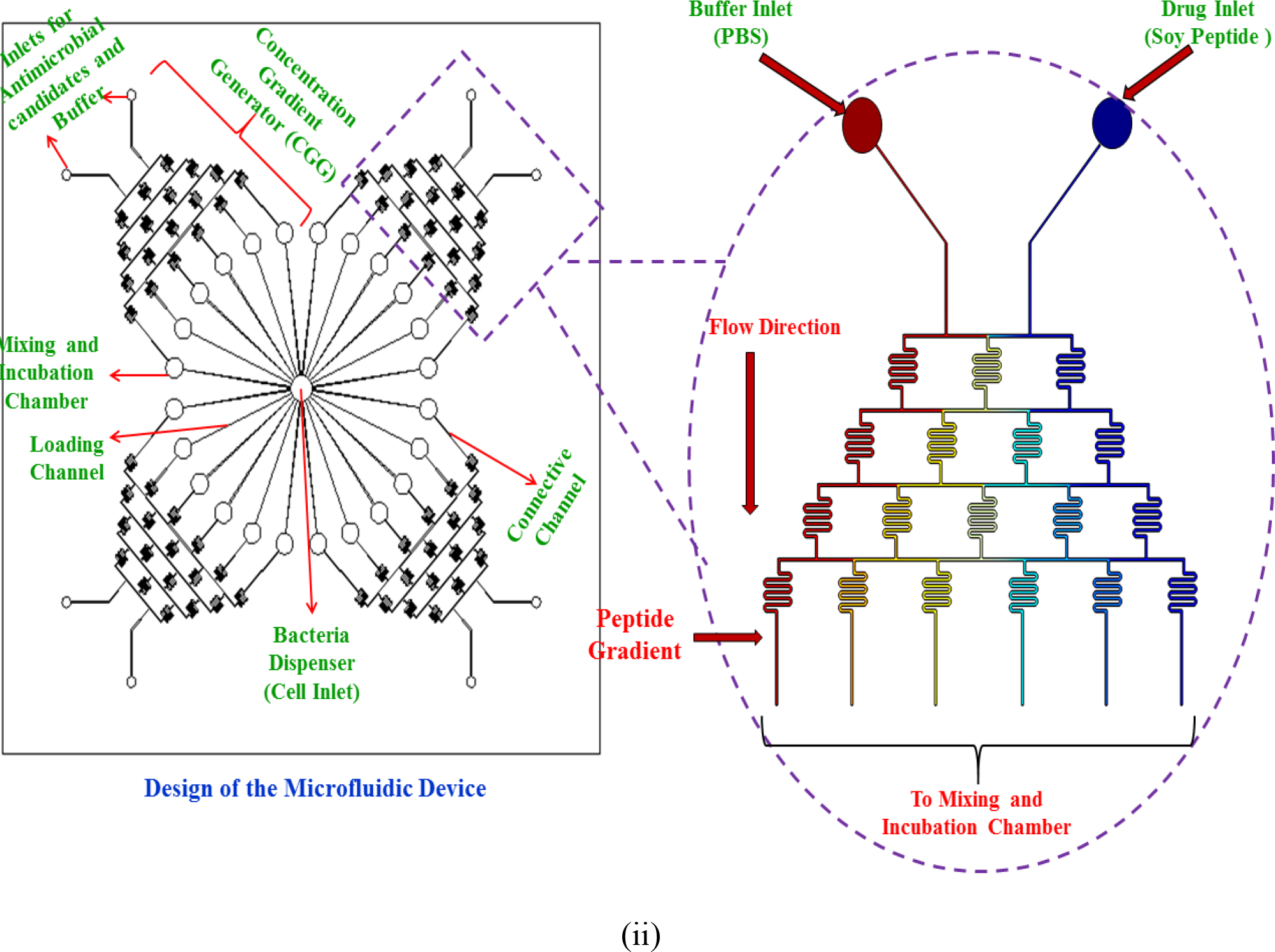
Design of the 3D microfluidic high-throughput drug screening platform: (i) three layers of the microfluidic device and a magnified view of the serpentine-like channels in the concentration gradient generator (CGG) and bacterial dispenser with incubation chambers. (ii) The top view of the microfluidic platform and a magnified view of the complete CGG with gradient generated by the soy peptide and buffer solution at the end of the generator.

### 2.6. Characterization of the Microfluidic Device

The fabricated 3D microfluidic device was characterized using fluorescein dye (Sigma Aldrich, Canada). This test was performed to ensure that the CGG generated steadily or linearly increasing concentrations at the connective channels. A 5 mM stock solution of fluorescein was prepared in water. A 5 µM solution was prepared from this stock and injected in one of the inlets of a CGG along with water at the other inlet of CGG. Syringe pumps were used for controlled injection (20 µl/h) of water and fluorescein dye into the CGG (Chemyx Fusion Touch, USA) through a 1 mm syringe and 0.3 mm needle tips (Becton, Dickinson and Company, USA). Images were taken using an inverted Nikon Eclipse Ti microscope with a green filter. Images were processed using ImageJ software (NIH, USA).

### 2.7. Cell Seeding into the Microfluidic Device

*P. aeruginosa* and *L. monocytogenes* treated with IKAFKEATKVDKVVVLWTA (long chain) peptide were used for all microfluidic experiments. The cell suspensions were prepared from overnight cultures grown at 37°C in their respective broth solutions. One ml of this culture was centrifuged (SciLogex D3024, USA) at 3000 rpm for 3 minutes. The supernatant was removed, and the pellet was suspended in sterile PBS. This was then diluted to a 0.5 McFarland Standard and used for further experimentation to ensure that cells would be poised to grow once inoculated. The cells were injected into the bacterial dispensing chamber at the center of the device. Once the loading channels and 24 chambers started to fill with bacteria, soy peptide at a concentration of 4.6 µM, 37.2 µM, 74.4 µM, and 298 µM was injected into the inlets of 4 CGGs, along with PBS buffer, using 1 mm syringes with 0.3 mm needle tips (Becton, Dickinson and Company, USA). The flow rate (20 µl/h) was controlled using syringe pumps (Chemyx Fusion Touch 200 and 400, USA; Harvard Apparatus Pump 11 Elite, USA). The temperature of the device was maintained at 37°C using a temperature controller stage. Once the concentration gradients start to fill and the 24 wells were filled with the peptide and bacterial mixture, the syringe pumps were turned off and the bacteria were incubated in the device.

### 2.8. RAMAN Spectroscopy Analysis

RAMAN Spectroscopy was used in identifying *P. aeruginosa* and *L. monocytogenes* before and after various treatments with long chain peptides. This was done to emphasize that the spectral signature of the bacteria and peptide mixture at different dosage levels could be used to validate the high throughput drug screening microfluidic device and show the efficacy of peptides against bacteria. *P. aeruginosa* and *L. monocytogenes* were incubated at a 1:1 (v/v) ratio of long chain peptide (IKAFKEATKVDKVVVLWTA) for 4 h at 37 ␥C to obtain final peptide concentrations of 4.6 µM, 37.2 µM, 74.4 µM, 100 µM, and 298 µM. RAMAN spectra was obtained using a 785 nm RAMAN spectrometer (Snowy Range Instruments, WY, USA) for control and all treatments on a surface enhanced (SERS) gold substrate (5 mm X 5 mm; Nanova Inc, USA). Spectra obtained on glass substrates (without SERS substrate) did not show any visually significant peaks or differences compared to the peptide treated bacterial samples (data not shown). Ampicillin was used as a control in this experiment.

### 2.9. Time Lapsed Microscopy Analysis

A 0.5 McFarland standard of *L. monocytogenes* (in logarithmic growth phase) was mixed with the long chain soy peptide to obtain final concentrations of 37.2 µM and 200 µM of soy peptide in microfuge tubes. A 10 µl of this mixture was added to the well in a microfluidic device and immediately placed under a Nikon Eclipse Ti inverted microscope (Nikon Instruments Inc., Melville, NY). A total 10 µl of the solution ensured that the individual bacterial cells were exposed to sufficient volumes of media. Images were captured focusing on the same region of interest over a 3 h time period with a 40x objective lens in the phase contrast 1 mode, equipped with the NCB and D filters. This experiment was performed to visualize the antimicrobial effect of the long chain soy peptide on bacterial cells. All experiments were performed in triplicate and analyzed using IBM SPSS Statistics 22.

## 3. Results and Discussion

### 3.1. Choice of Peptides for this Study

After a detailed literature review, we determined that the therapeutic potential and antimicrobial properties of soy based peptides have not been widely studied. Therefore, two peptides of soy origin (PGTAVFK and IKAFKEATKVDKVVVLWTA) were chosen for this study based on three published papers that investigated: 1) the antimicrobial activity of PGTAVFK against *Escherichia coli* and *Staphylococcus aureus*, 2) the activity of fragment (213-231) of the enzyme D-myo-inositol-3-phosphate synthase from soybean (GenBank accession number: ABM17058.1) against Asian rust spores, and 3) the activity of myoinositol as a termicide [3, 28, 39].

Bacterial protein phosphorylation has been recently identified as a promising target for the development of new antibacterial compounds [5]. D-myo-inositol 3-phosphate synthase is an enzyme that has the ability to catalyze the production of phospholipids during salvage. Inositol is a metabolic sensor for numerous signal transduction pathways that specifically has the ability to disrupt cellular processes during the unfolded protein response [17]. The sequence of fragment (213-231) from D-myo-inositol-3-phosphate synthase is IKAFKEATKVDKVVVLWTA [3].[3]. Therefore PGTAVFK and IKAFKEATKVDKVVVLWTA were chosen as antimicrobial peptide candidates for this study.

### 3.2. Antimicrobial Testing Using Microdilution Assay

Microdilution assays were performed to examine the antimicrobial efficacy of both short chain (PGTAVFK) and long chain (IKAFKEATKVDKVVVLWTA) soy peptides against *P. aeruginosa* and *L. monocytogenes*. Figure 2 (PGTAVFK) and Figure 3 (IKAFKEATKVDKVVVLWTA) show the effects of these two soy peptides on the optical density of bacterial cultures, which reflects their antimicrobial capacities (ability to stop/slow growth). PGTAVFK did not significantly affect *P. aeruginosa* culture density, but it had an inhibitory effect on *L. monocytogenes* at a concentration above 625 µM. IKAFKEATKVDKVVVLWTA, on the other hand, was effective against both *P. aeruginosa* and *L. monocytogenes* at concentrations above 37.2 µM. It is apparent from our results that both antimicrobial peptides can affect the optical density of the organisms, which represents the concentration of the suspended organisms and the sum total of growth and cell death. Therefore, we cannot speculate whether the major modes of actions for these peptides are bactericidal versus bacteriostatic; future studies will explore the precise mechanisms of action.

**Figure 2.**
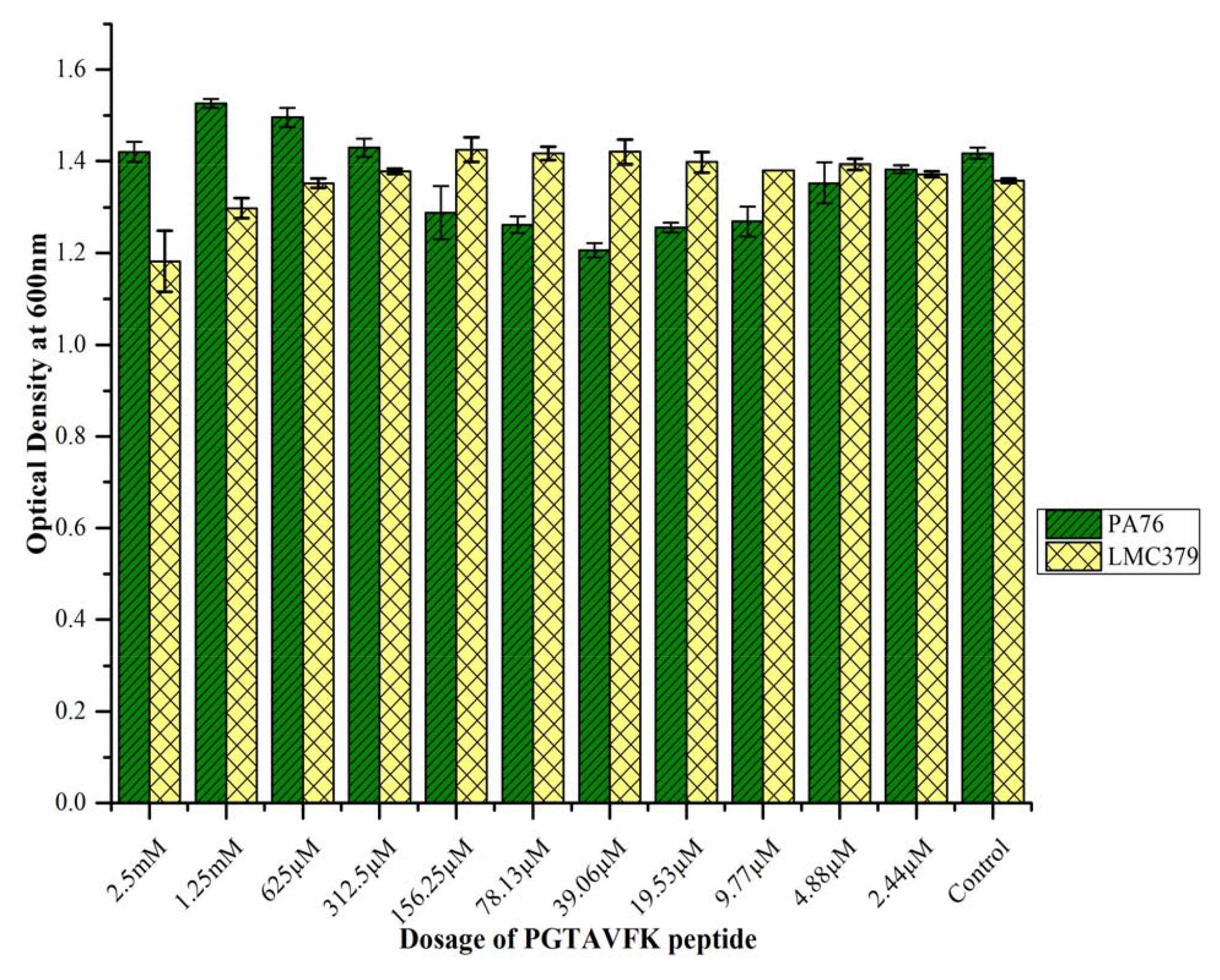
Antimicrobial activity of the soy peptide PGTAVFK against *P. aeruginosa* PA76 and *L. monocytogenes* C379. The soy peptide did not inhibit the bacterial growth of *P. aeruginosa* significantly but had an inhibitory effect on *L. monocytogenes* at concentrations above 625 µM.

**Figure 3.**
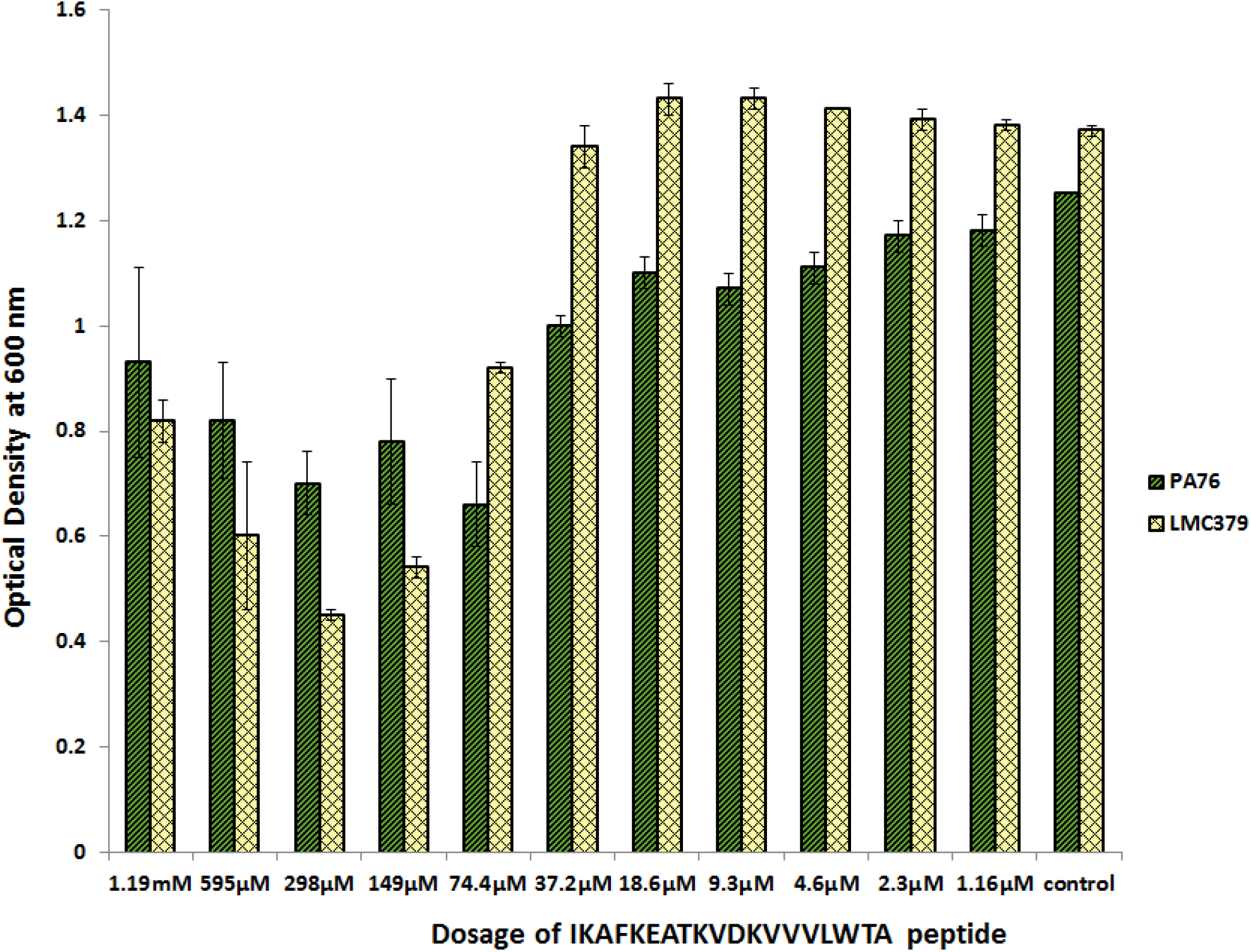
Antimicrobial activity of the soy peptide IKAFKEATKVDKVVVLWTA against *P. aeruginosa* and *L. monocytogenes*. The soy peptide inhibited both strains of bacteria at a concentration as low as 37.2 µM.

The difference in antimicrobial efficacy between the short and long chain peptides could be attributed to the structure and length of the sequences (7 amino acid sequence: PGTAVFK; 19 amino acid sequence: IKAFKEATKVDKVVVLWTA). As discussed previously, long chain peptides are generally associated with high antibacterial activity. The difference in the net positive charges of the two peptides could also have contributed to differences in their antibacterial activity. The activity of AMPs could be affected by the concentrations of cations and anions present in their environment [12]. Therefore, the combination of amino acids in the sequences of peptides could have affected their antibacterial efficacy.

It has been demonstrated that some cations neutralize repulsive forces of adjacent lipopolysaccharide (LPS) in the outer membrane of Gram-negative bacteria, leading to a tight and cross-linked outer membrane that protects itself from hydrophobic surfaces and AMPs [26]. LPS moieties differ in composition and length among bacterial species, which could possibly be a reason why the long chain peptide was able to act effectively on *P. aeruginosa*, while the short chain peptide was largely inactive [26]. This could also be the reason why PGTAVFK was not able to significantly inhibit *P. aeruginosa* but was able to inhibit *L. monocytogenes*, a Gram-positive bacterium. Once AMPs pass through the LPS in outer membrane, there is an inner membrane comprised of phospholipids on both sides of the bilayer. Therefore, a peptide that could enter the outer membrane need not necessarily interact efficiently with the inner membrane. It has been reported that peptides form aggregates in solution, which makes it difficult for them to enter the outer membrane and reach the target, which is the cytoplasmic membrane [26]. However, no aggregates were visually observed in either prepared peptide solution used in our experiments. LPS forms aggregates in solution that are biologically active and toxic. Therefore, dissociation of these aggregates is required to neutralize and inactivate LPS. These could be some plausible reasons for the observed differential sensitivity.

### 3.3. Characterization of our High Throughput 3D Microfluidic Platform

Characterization of the 3D microfluidic device was performed to show that this device could be used for multiplexing, parallel concentration gradient generation, and drug screening applying various concentrations simultaneously. Moreover, the use of PDMS helps in creating an inexpensive, biocompatible, and transparent environment that allows the use of this device in further quantification through optical and RAMAN imaging.

Under continuous flow conditions, the 3D high throughput device was characterized using 5 µM fluorescein dye. Figure 4 shows the concentration gradient formed in the six channels in terms of fluorescence intensity. Diffusion of fluorescein into the PDMS layer forms concentration gradients. Images of fluorescence in all six channels were obtained using Nikon Ti inverted microscope, and the intensity of fluorescence was then measured from 6 channels and normalized. We found that the fluorescence intensity increases with increasing channel number. Formation of air bubbles in the serpentine-like pathways of the microfluidic platform may block fluid flow causing non-linearity, which can be avoided by prober bonding of the PDMS with the glass slide and an accurate device design chip. It is quite important that the thickness of PDMS layers is uniform for a proper concentration gradient generation. Our group and others have successfully demonstrated that it is possible to generate a linear concentration of drug molecules using a microfluidic device of similar design; results from this study are consistent with these previous studies [7, 16].

**Figure 4.**
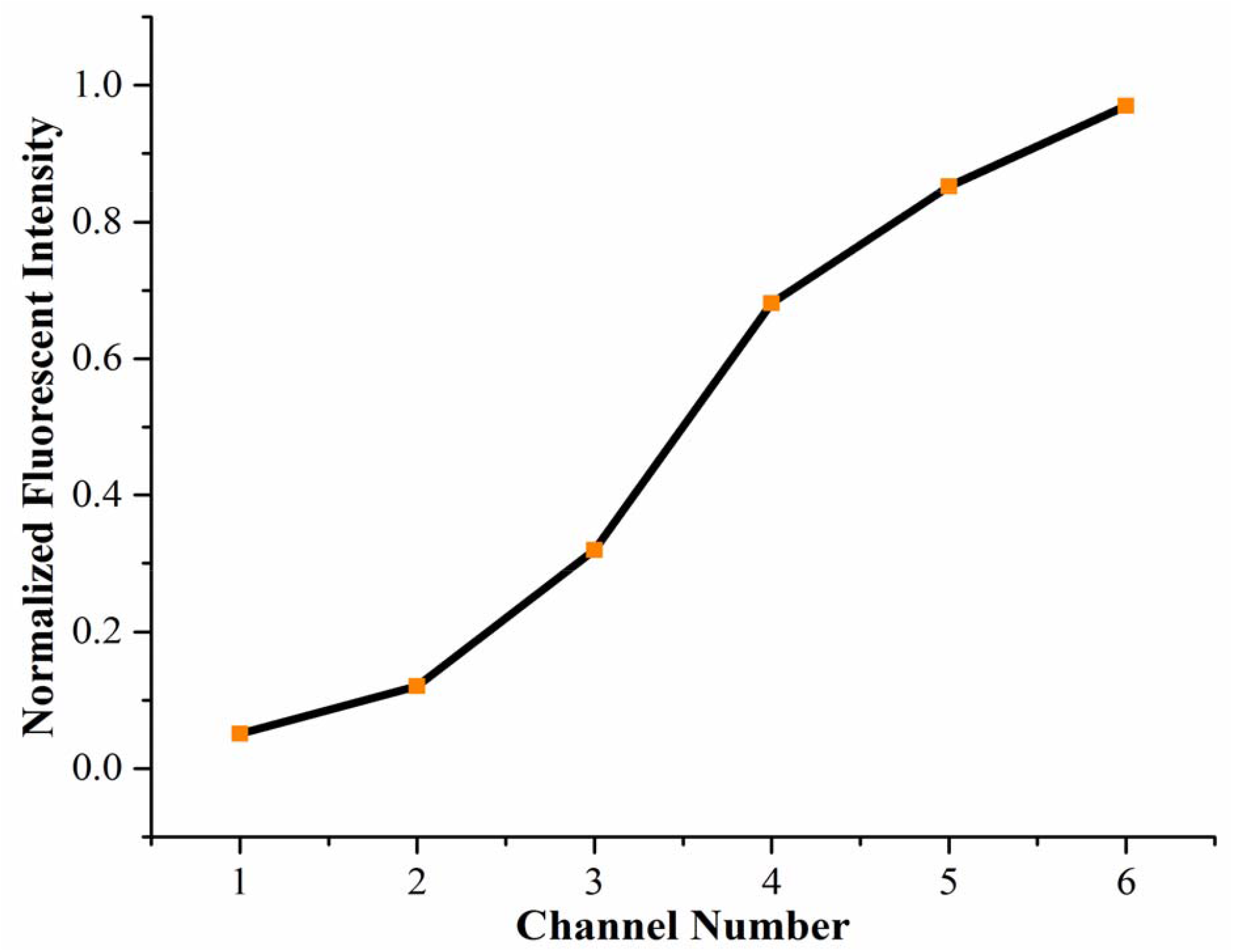
Characterization of the 3D microfluidic high throughput drug screening device with 5 µM of fluorescein and buffer at the inlets of device at a flow rate of 20 µl h^-1^. The relationship between the normalized fluorescent intensity and the channel number shows that intensity increases with increasing channel number implying that the concentration gradient generator of the device (CGG) produces linear concentrations of fluorescein in the device.

### 3.4. RAMAN Spectra Analysis

RAMAN spectroscopy is a preferred tool for identifying and detecting pathogens for point-of-care diagnostics applications [2, 31]. Here, it was used in conjunction with the 3D microfluidic platform to validate that by reducing the sample volume to few microliters in a microfluidic device, the incubation time for detection could be reduced drastically to identify bacterial species. Owing to the microliter volume size of the peptide solutions in each well of the microfluidic incubation chamber and the enhanced surface to volume ratio, there is a rapid kinetic reaction between the peptide and the individual bacterial cells, leading to a decreased incubation time in comparison with the milliliter volumes of the conventional assays. We recorded the SERS spectra at an excitation wavelength of 785 nm under various dosage levels of long chain peptide (IKAFKEATKVDKVVVLWTA), treating both *P. aeruginosa* and *L. monocytogenes* for 4 h at 37° (Figure 5). We found that for the *P. aeruginosa* control, the RAMAN peaks were visible at about 823, 1270, 1362, 1490, and 1510 cm^-1^ (highlighted and circled in Figure 5 (i)). For the *L. monocytogenes* control, peaks were visible at about 716, 821, 1022, 1423, and 1454 cm^-1^ (boxed and circled in Figure 5 (ii)). In both cases, these significant peaks could not be observed in spectral curves of the treated samples, which suggest that the peptides possess antimicrobial activity.

In an attempt to identify spectra of biomolecules found in microorganisms using an excitation wavelength of 785 nm, DNA and RNA bases were observed to dominate the region between 600 and 800 cm^-1^; proteins in the form of amide I and III ranged from 1300 to 1655 cm^-1^, fatty acids fell in the range of 1300-1440 cm^-1^, and saccharides were between 1000 and 1200 cm^-1^ and1300 and 1500 cm^-1^ [6]. The speculated molecular characterization of spectral peaks observed in this study, with reference to the work of De Gelder *et al* [6], is shown in Table 1. The spectral peaks observed for *P. aeruginosa* could be attributed to the presence of various forms of fatty acids and DNA and RNA bases, while in *L. monocytogenes*, the peaks could be attributed to the presence of fatty acids, saccharides, and amino acids.

**Figure 5.**
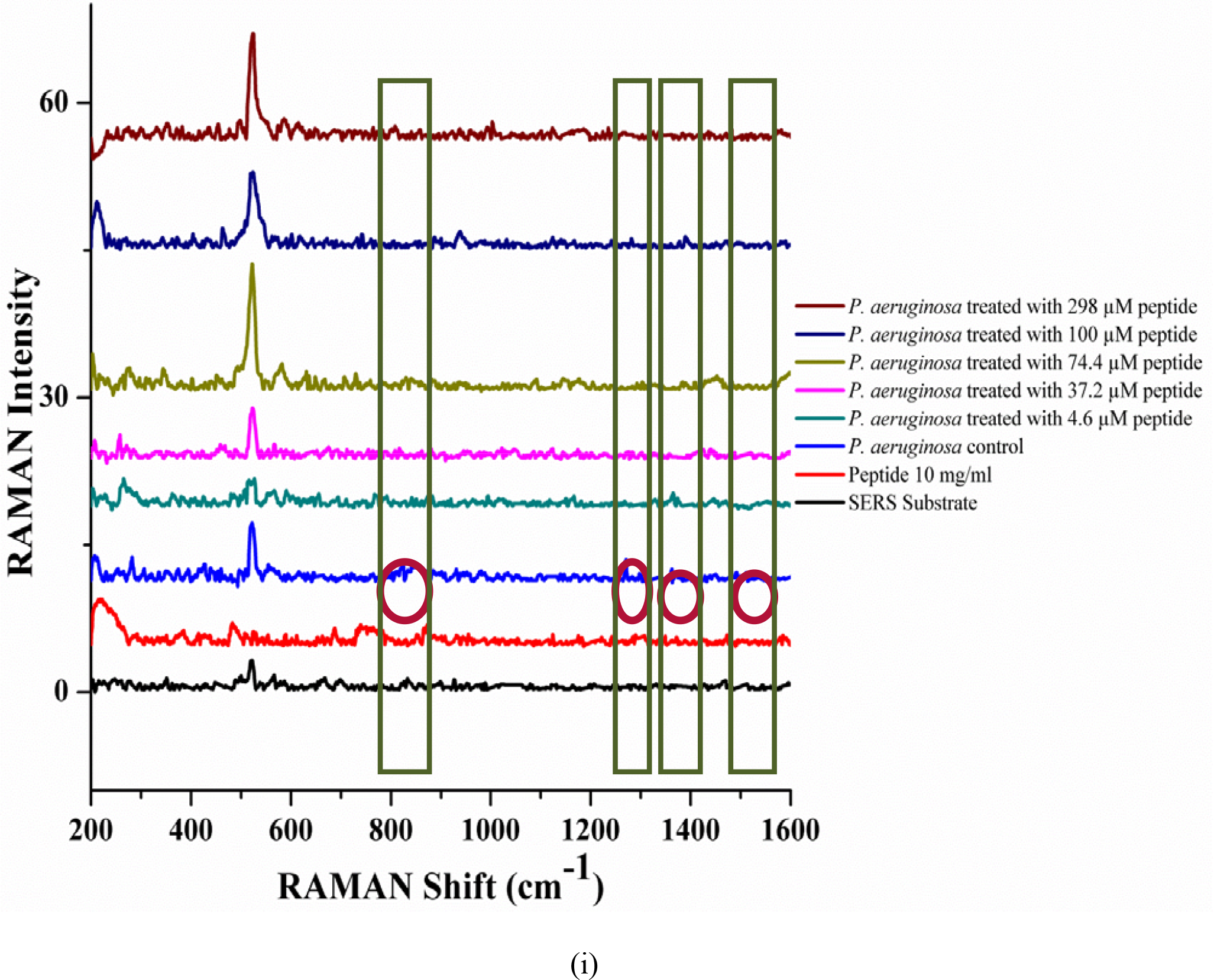
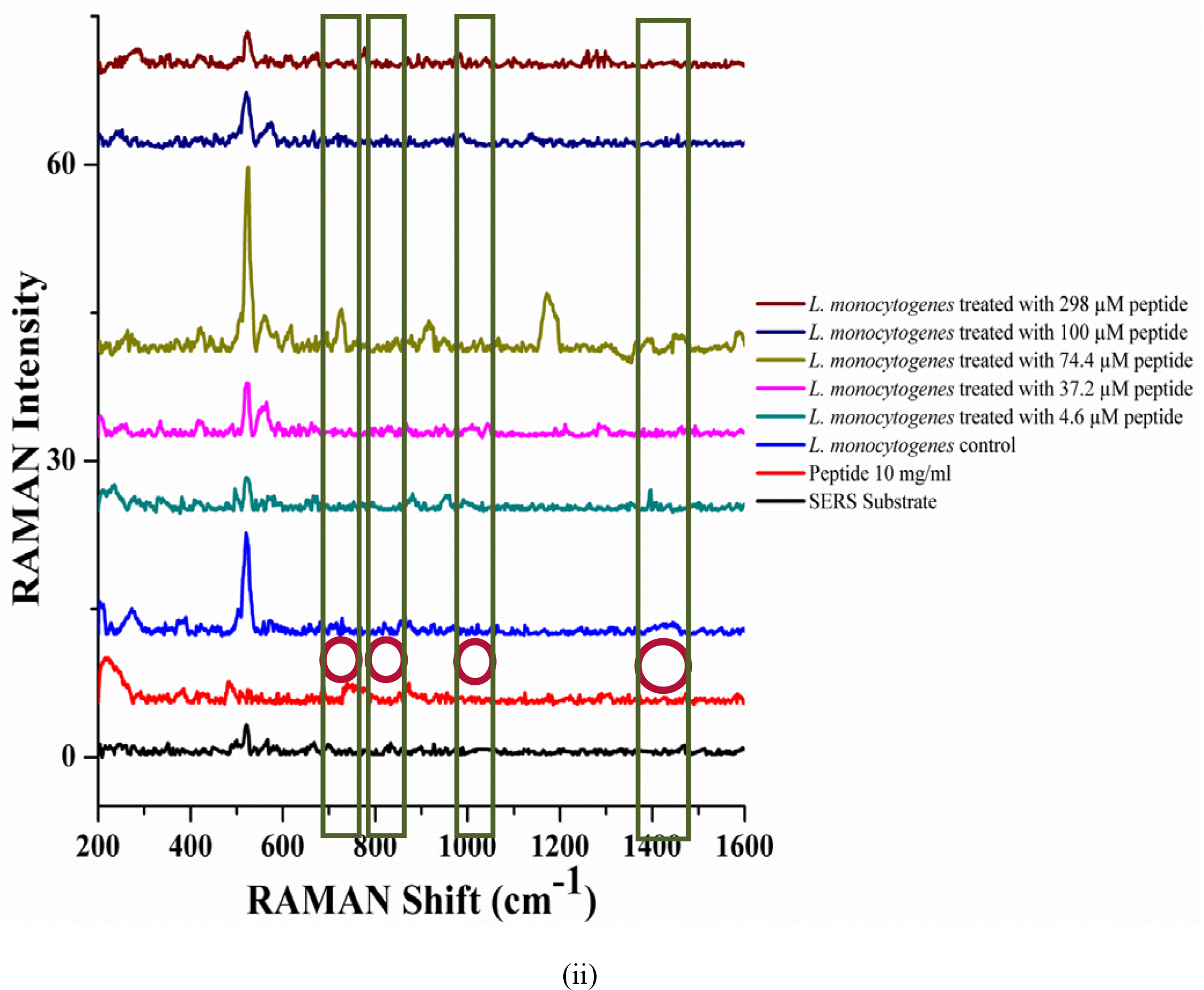
**Surface Enhanced RAMAN Spectra (**SERS) obtained for (i) *P. aeruginosa* and (ii) *L. monocytogenes* with and without 4.6, 37.2, 74.4, 100, and 298 µM long chain peptide treatments. Shifts in the control spectra are circled and their comparisons with different treatments are boxed.

**Table 1.**
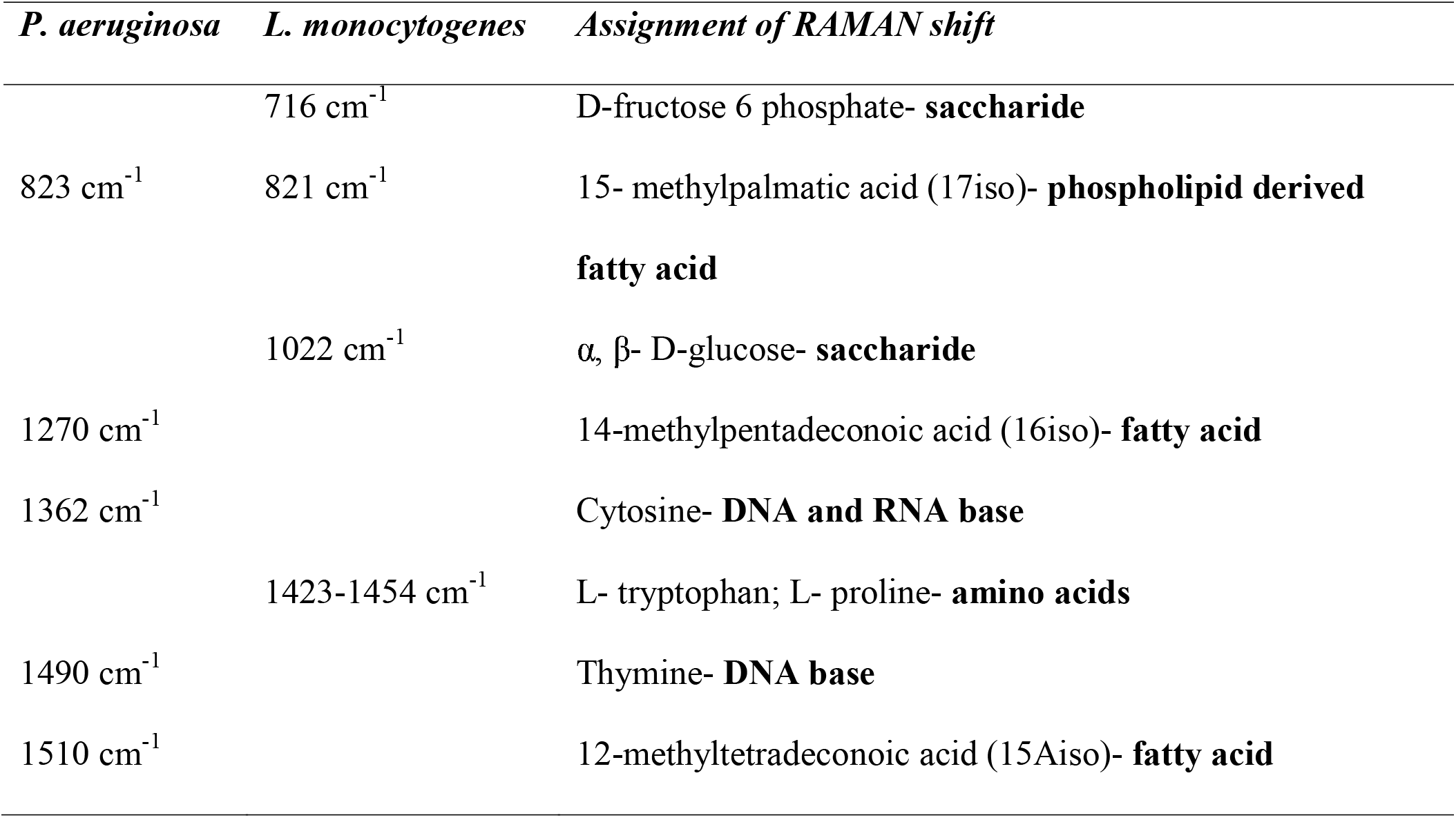
Speculative assignment of peaks from the SERS spectra of *P. aeruginosa* and *L. monocytogenes* using a 785 nm excitation wavelength based on the characterization reported by De Gelder *et al* [6].

Conventional techniques employed to identify bacteria are usually slow, cannot be used at field sites, and require a technical personnel to operate. Therefore, SERS is preferred due to the possibility of obtaining high vibrational fingerprint information at an increased sensitivity (to even single molecule) and the possibility of using it in aqueous conditions with little or no interference [8]. The SERS substrate enhances the RAMAN intensity of samples due to the surface roughness caused by the presence of metal nanoparticles (gold). Molecules in the samples are adsorbed onto these surfaces, which increases the strength of detection.

When RAMAN spectra was obtained from the samples on a glass substrate, the spectra of both bacteria, including treatments, looked similar without any distinguishable shifts (data not shown), which could be attributed to low sensitivity of the method. By using a SERS substrate, the spectral intensity improved, but the RAMAN shifts were still weak. It was speculated that the incubation time and bacterial cell density have a direct correlation. If the bacterial cell density is less, the incubation time could also be reduced and so forth. In these experiments, a 0.5 McFarland standard cell density was used (OD at 600 nm was between 0.08 and 0.1). Therefore the amount of incubation time required to elicit visible strong changes in biochemical signatures at this cell density could be longer than 4 h.

Sockalingum *et al* [38] used different ways of mixing metal colloids with bacteria and observed SERS spectra using 632 nm excitation. It was found that the spectra of *E. coli* differed from that of *P. aeruginosa* by observing peaks of constituents, such as fatty acids, amino acids, proteins, and nucleic acids. Using a 784 nm excitation, Guzelian *et al* [15] claimed that there were different spectra for *P. aeruginosa, L. monocytogenes, Bacillus subtilis*, and *Bacillus cereus* when placing bacteria on electrochemically roughened gold SERS strips. At 514, 633, and 780 nm, Efrima & Zeiri [8] have also used the SERS technique to identify molecular information for bacteria. Therefore, reproducibility and selectivity remain a drawback. However, when controlling experimental conditions, Efrima & Zeiri [8] demonstrated that these drawbacks can be overcome.

The low sensitivity of the tabletop RAMAN instrument itself could be another reason for the weak signatures obtained in our study. In order to further discriminate the differences in the spectral data, discriminant analysis was performed to show that the function was able to classify 100% of *P. aeruginosa* cells and 85.9% of *L. monocytogenes* cells correctly.

### 3.5. Time Lapsed Image Analysis

The 3D high throughput microfluidic platform was used in conjunction with optical microscopy to image the morphology of *L. monocytogenes* treated with 37.2 and 200 µM concentrations of IKAFKEATKVDKVVVLWTA peptide at 0, 1, 2, and 3 h. This experiment was performed to visualize the effect of peptide on *L. monocytogenes* cells. Figure 6 shows the time lapsed images of *L. monocytogenes* treated with 37.2 and 200 µM peptide taken at 40X magnification over a period of 3 h. Aggregation in the peptide-treated sample is significantly lower than in the case of untreated cells The motility of the cells was restricted when comparing 2 and 3 h treatments with 200 µM of peptide used on *L. monocytogenes*. The effect of antimicrobial soy peptide was observed to inhibit flocculation and reduce the motility of bacterial cells.

**Figure 6.**
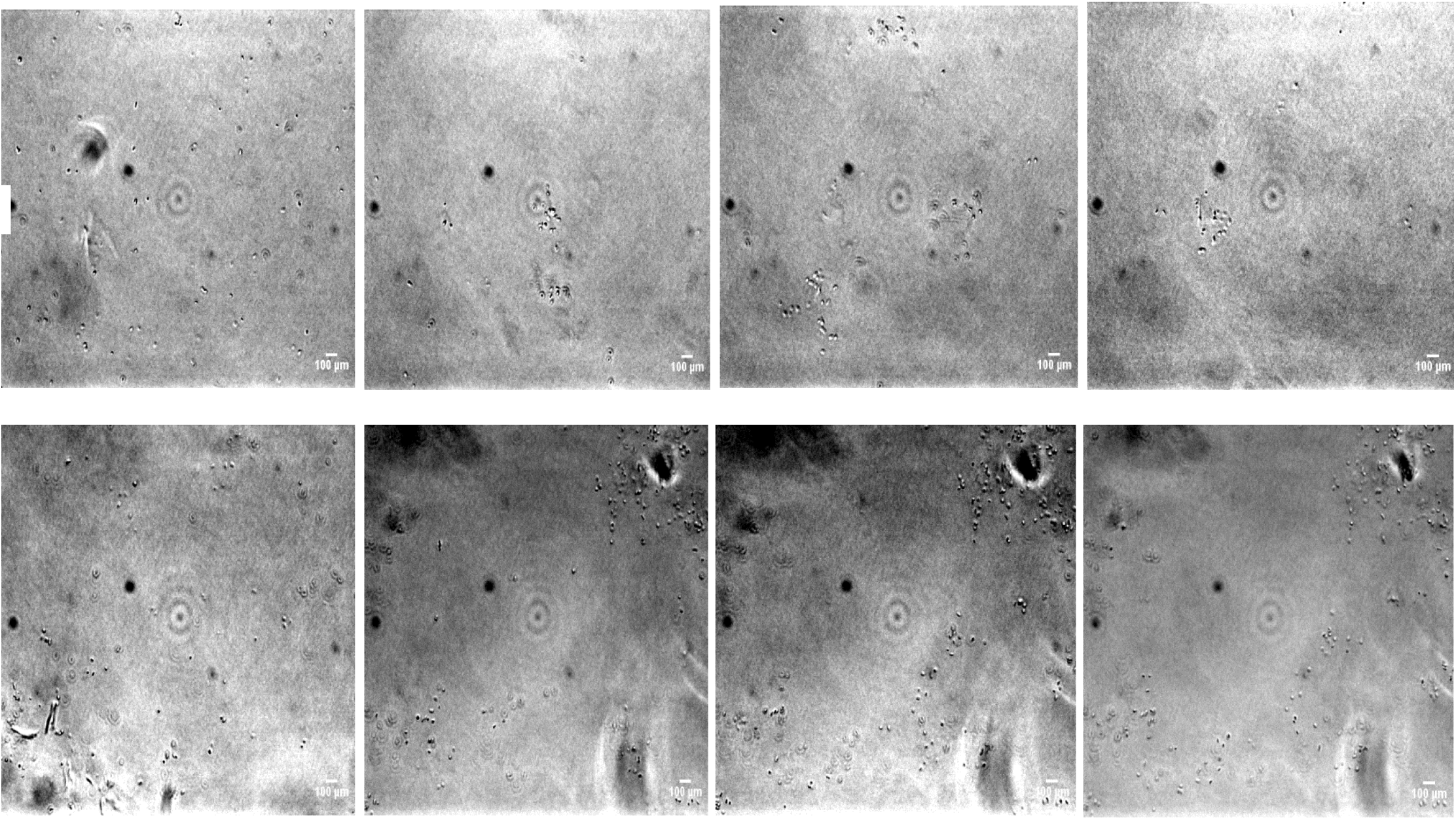
Time lapsed images of *L. monocytogenes* treated with 37.2 µM and 200 µM of long chain soy peptide in the microfluidic device at 0, 1, 2, and 3 h taken with the 40x lens of a Nikon Eclipse Ti inverted microscope.

## 4. Conclusions

Antimicrobial peptides are a novel class of bio-pharmaceuticals that possess significant therapeutic potential. The results of this study indicate that soy peptides PGTAVFK and IKAFKEATKVDKVVVLWTA could possibly be used in the development of plant based, biocompatible, and biodegradable alternative antimicrobial agents or antibiotics for use in the food industry and medical field.

There is a dire need to develop and exploit high throughput techniques for drug analysis andantimicrobial/antibiotic screening in the pharmaceutical industry. A high throughput, sensitive, and selective tool would drastically reduce the timelines for antimicrobial candidate discovery and development within the pharmaceutical industry. Because of the ability to integrate 3D microfluidic high-throughput drug screening platform with microscopy and spectroscopy, rapid evaluation of antibiofilm candidates would help to prevent false positive responses and will ultimately streamline preclinical trials.

## Acknowledgements

The authors sincerely thank the Natural Sciences and Engineering Research Council of Canada, Ontario Ministry of Research and Innovation, and the Ontario Ministry of Agriculture and Food for funding this study.

